# A note on discretising Keyfitz entropy

**DOI:** 10.1101/2022.09.05.506601

**Authors:** Charlotte de Vries, Connor Bernard, Roberto Salguero-Gómez

## Abstract

Keyfitz’ entropy is a widely used metric to quantify the shape of survivorship of populations, from plants, to animals, and microbes. Keyfitz’ entropy values < 1 correspond to life histories with an increasing mortality rate with age (i.e., actuarial senescence), whereas values > 1 correspond to species with a decreasing mortality rate with age (negative senescence), and a Keyfitz entropy of exactly 1 corresponds to a constant mortality rate with age. Keyfitz’ entropy was originally defined using a continuous-time model, and has since been discretised to facilitate its calculation from discrete-time demographic data. In this short note, we show that the previously used discretisation of the continuous-time metric does not preserve the relationship with increasing, decreasing, or constant mortality rates. To resolve this discrepancy, we propose a new discrete-time formula for Keyfitz’ entropy for age-classified life histories. We show that this new method of discretisation preserves the relationship with increasing, decreasing, or constant mortality rates. We analyse the relationship between the original and the new discretisation, and we find that the existing metric tends to underestimate Keyfitz’ entropy for both short-lived species and long-lived species, thereby introducing a consistent bias. To conclude, to avoid biases when classifying life histories as (non-)senescent, we suggest researchers use either the new metric proposed here, or one of the many previously suggested survivorship shape metrics applicable to discrete-time demographic data such as Gini coefficient or Hayley’s median.

## 1 Introduction

Actuarial senescence is defined as the increased risk of dying as an individual gets older (Medawar, 1952). Getting older cannot be avoided in that it is a natural consequence of surviving, but some species seem to be able to avoid senescing (Vaupel *et al*., 2004; Jones *et al*., 2014; Roper *et al*., 2021). It is tempting to assume long-lived organisms senesce less than short-lived organisms, but of course it is possible for an organism to have a constant but high mortality rate over its entire lifespan, and thus be (relatively) short-lived and negligibly senescent (Péron *et al*., 2019). For example, Baudisch (2011) compare 10 animal species and find that robins (*Erithacus rubecula*) rank as having the shortest life expectancy while at the same time having the lowest pace of life, that is, the least senescent survivorship curve (out of this admittedly small sample of 10 animal species). Likewise, an organism can have an increasing but relatively low mortality rate over its entire lifespan, and thus be long-lived and senescent. For example, bamboo (Phyllostachys) stands rapidly die following a period of relatively low mortality lasting 60-100 years (Janzen, 1976; Finch & Rose, 1995). Similarly, long-lived semelparous plants such as long-lived Puya raimondii (living up to 150 years; Finch (1998)) and Agave americana (which often live decades; Harper & White (1974)) show delayed, but rapid declines in vitality with age.

To disentangle these two dimensions of ageing, namely longevity and the rate of senescence, demographers typically distinguish between the pace of ageing and the shape of ageing, respectively (Keyfitz, 1968, 1977; Baudisch, 2011). The *pace* of life is often quantified through demographic metrics such as mean life expectancy, reproductive window, or generation time, which tend to be highly correlated. The pace of ageing behaves intuitively: it is high for short-lived organisms and low for long-lived organisms. The *shape* of ageing, on the other hand, is determined by the time-standardized shape of the mortality or survival curve. The goal of shape metrics is to classify survival curves by whether the mortality rate mostly increases or decreases with (standardized) time (respectively, scenescent versus negative senescent curves), see Figure 1 for some examples of different survivorship curve shapes.

**Figure 1:**
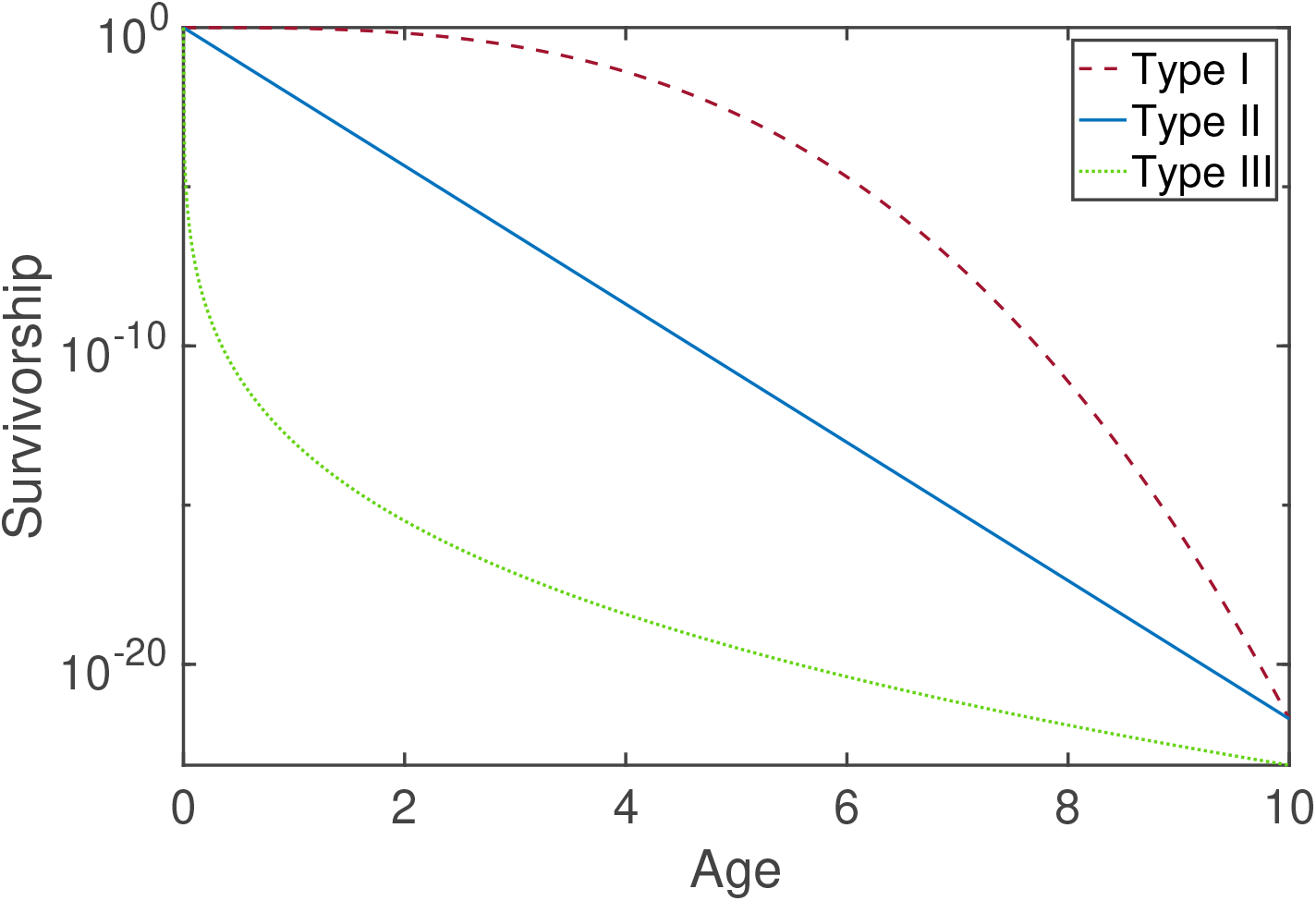
Three example survivorship functions of the three different types (type I: senescence; type II: constant mortality; type III, negative senescence). The two Keyfitz entropy measures given by equations (5) and (10)) are calculated for these three curves using two different widths of the age classes, Δ*t*, are given below in Table 1. That is, we discretized the survivorship curves shown in the figure using two different sizes of discrete intervals, Δ*t* = 0.01 and Δ*t* = 1.

Keyfitz’ entropy is one of the metrics that has been proposed to quantify to shape of ageing (Keyfitz, 1977; Wrycza *et al*., 2015). Keyfitz’ entropy was originally identified as a dimensionless measure of the elasticity of lifespan to a uniform change in age-specific mortality (Leser, 1955). Population entropy was later re-derived and popularised by Keyfitz (1977). A similar measure was introduced through independent proofs by Demetrius, which helped attract interest to the measure (Demetrius, 1974, 1978). Demetrius (1978) noted the potential use of Keyftiz’ entropy for the classification of survivorship curves, pointing out that it has the useful property that *H* = 1 corresponds to a constant mortality rate (type II curve), *H* < 1 corresponds to mortality increasing with age (type I curve), and *H* > 1 corresponds to mortality decreasing with age (type III curve, see figure 1 for an example of all three curve types).

Salguero-Gómez *et al*. (2016) introduced a discretized version of Keyfitz entropy that interchanges the integral for summation which has been subsequently used in a number of publications (Salguero-Gómez, 2017; Beckman *et al*., 2018; Capdevila *et al*., 2020; Bernard *et al*., 2020). In this short note, we show that this approach to discretise Keyfitz entropy does not fully capture the expected relationship with increasing, decreasing and constant mortality rates of the continuous-time metric. To resolve this discrepancy, we introduce a different discrete-time version of Keyfitz’ entropy based on previous work on matrix formulas for life disparity (Vaupel & Canudas-Romo, 2003; Keyfitz & Caswell, 2005; Caswell, 2013; Caswell *et al*., 2018). We show that this alternative discretisation does preserve the expected relationship between Keyfitz’ entropy, and whether/how mortality rates change with age in age-structured matrix population models. We then analyse the relationship between the two Keyfitz’ metrics to test if any consistent biases might exist. We evaluate this relationship empirically using animal and plant matrix population models, as discussed in more detail below. We find that the two metrics classify survivorship with a similar profile across values, with a consistent, strong trend toward underestimating entropy values (suggesting stronger senescence) using the original entropy metric. That is, curves classified as (weakly) negatively senescent by the new entropy metric are likely to be incorrectly classified as senescent by the original entropy metric, and constant mortality curves are regularly incorrectly classified as senescent curves by the original entropy metric. In addition, life expectancy strongly correlates with deviation between the two entropy metrics, with long-lived species showing little bias and short-lived species showing a stronger bias in the original Keyfitz metric.

## 2 Methods: Keyfitz entropy

### 2.1 Continuous-time formulas for Keyfitz entropy

Keyfitz (1977) (or the recent edition, Keyfitz & Caswell (2005)) defines the measure *H* of a survival function *l*(*a*) at age *a* as

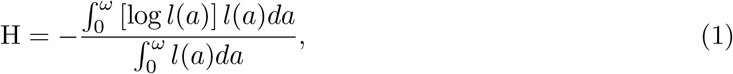

and they note that *H* has been called entropy of information in other contexts (section 4.3 of Keyfitz & Caswell (2005)). *H*, since then generally known as Keyfitz’ entropy, as defined above is a weighted average of the logarithm of survival, where the weights reflect normalised survivorship. The use of entropies and information theory has a long history in ecology and evolution (e.g., Shannon biodiversity indices, Kullback-Leibler and Jenzen-Shannon divergence, and Max Ent distribution modelling). Note, however, that Keyfitz’ entropy does not integrate to one and is therefore not an entropy in the strictest sense (Shannon & Weaver, 1949).

Keyfitz & Caswell (2005) derived the formula by calculating the effect of a proportional change in age-specific mortality on the life expectancy at birth, and the measure therefore relates to the similarity of mortality across age-classes. As a result, Keyfitz’ entropy is also a measure of the concavity of the survivorship curve. Goldman & Lord (1986), Vaupel (1986), and recently Vaupel & Canudas-Romo (2003) showed that Keyfitz entropy can be decomposed into two constituent measures that demographers use: 1. Life disparity, *e*^†^, which is defined as the average remaining life expectancy at the ages when death occurs, and measures the number of life years lost due to death 2. Average lifespan of an individual at the time of birth, *e*^0^, which is calculated by integrating the survival density function. The ratio of the aforementioned indices represents an equivalent formulation of Keyfitz’ entropy:

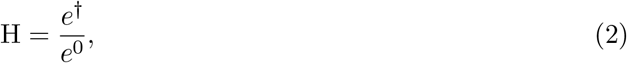

whereby

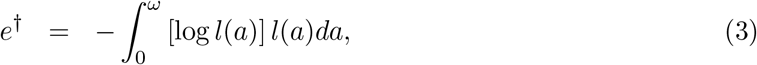

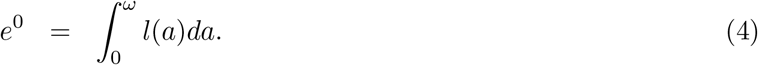

### 2.2 Discretisation of Keyfitz entropy

Salguero-Gómez *et al*. (2016) introduced a discretised version of Keyfitz entropy,

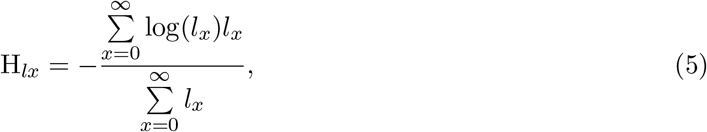

see Table S2 from the Supplementary Materials of Salguero-Gómez *et al*. (2016). A few lines of algebra will show that the continuous time version of Keyfitz’ entropy, equation (1) is equal to 1 when the mortality rate, *μ*, is constant such that *l*(*x*) = exp(−*μx*). The discretization in equation 5 (what we refer to as the original discret-time entropy measure) no longer has this property as can be seen from the Keyfitz entropies in Table 1, calculated for three example survivorship curves in Figure 1. In Supplementary Materials 1, we calculate the value of H_*lx*_ when mortality is constant and show that it is a function of *μ* which approximates one as mortality gets close to zero.

**Table 1:** Comparison of the two Keyfitz entropy discretisations for the curves in Figure 1 for a discrete time interval of Δ*t* = 0.01 and a discrete time interval of Δ*t* = 1. Keyfitz lx is given by equation (5), and Keyfitz N is given by equation (10).

| Entropy measure | Keyfitz lx |  | Keyfitz N |  |
| --- | --- | --- | --- | --- |
| | $\Delta t = 0.01$ | $\Delta t = 1$ | $\Delta t = 0.01$ | $\Delta t = 1$ |
| Type I | 0.33 | 0.30 | 0.33 | 0.44 |
| Type II | 0.96 | 0.75 | 1.00 | 1.00 |
| Type III | 1.85 | 0.76 | 1.94 | 1.18 |

### 2.3 An alternative discretisation of Keyfitz entropy

We propose an alternative discrete-time formula for Keyfitz’ entropy, derived from the definition of Keyfitz’ entropy as the ratio of life disparity to life expectancy at birth (which we refer to as the new discrete-time entropy measure). Starting from the fundamental matrix **N** (Caswell (2001), p. 112), life expectancy at birth is defined as

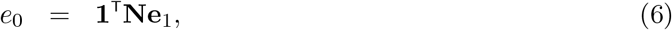

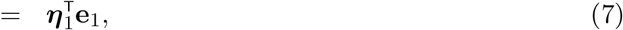

where ***η***_1_ is the vector of life expectancies at each age, **1**^⊤^ is a vector of ones, and **e**_1_ is a vector with zeros in all entries except the first entry which is one (see for example Caswell (2013) or Caswell *et al*. (2018)). Similarly, life disparity at birth can be calculated from the fundamental matrix **N** and the mortality matrix **M** as

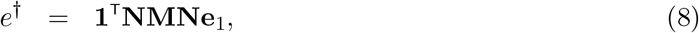

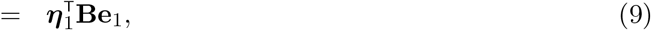

where the matrix **B** contains the distribution of age at death for an individual of each age, and the vector **Be**_1_ selects the distribution of age at death for a newborn individual, see also equation (57) in Caswell *et al*. (2018) or section 3 in Caswell (2013). Keyfitz’ entropy is then given by

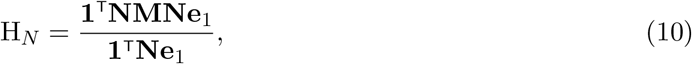

or equivalently by

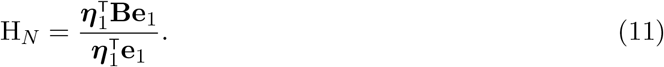

In Supplementary Materials 2, we show that equation (11) yields a value greater than 1 when mortality rate *μ* is a decreasing function of age, a value less than 1 when mortality is an increasing function of age, and exactly 1 when mortality is constant.

### 2.4 Comparing the metrics in real species using the COM(P)ADRE databases

To compare how the two discrete-time entropy measures perform in the context of biologically realistic models, we calculated entropies for age-structured matrix population models. We evaluated both animal and plant population models from the COMADRE (version 4.21.8.0) and COMPADRE (version 6.22.5.0) matrix population databases (both available from https://www.compadre-db.org). The empirical comparison included 401 species animal matrix population models and 34 species plant matrix population models.

We initially screened the matrix population models in COMPADRE and COMADRE based on their inclusion in previous publications that used Keyfitz’ entropy (Bernard *et al*., 2020). These models were selected based on duration of study, whether the population monitored was subject to experimental manipulation, among other criteria. Within the abovementioned subset, we selected unique records where duplicates existed based on maximizing study duration. We evaluated whether models were age or stage classified and removed any records that included NA values in reproductive elements or stage-specific survival values greater than unity.

For models from previous analyses that were stage-based (80 of 400 in COMADRE; 148 of 150 in COMPADRE), we converted them to age-based matrices. The stage-to-age transformation was based on life table projections of the stage matrix (Jones *et al*., 2022). We constructed Leslie matrices using survival vectors (px) and fertility vectors (mx) for ages within 90% of the starting population size based on the survivorship curve (lx). We compared higher and lower thresholds of the cohort size cut-off and found little variation in the number of viable age-specific analogues to stage matrices. Age-specific models converted from stage specific models were validated to see if the intrinsic population growth rate, reproductive rate, and generation time were consistent with those of the stage matrix that generate the life table from which the age-models were calculated. We only retained stage-based models where differences between stage and their age-equivalent matrices were within 5% of one another along the above demographic metrics. In most cases the differences were < 1%.

## 3 Results: bias of the existing metric correlates with longevity

To compare the original and new discrete-time entropy measures, in Table 1 we calculate the entropy using both measures for a few example survivorship curves shown in Figure 1. Both metrics change as the step size is changed. However, the new discrete-time entropy metric remains on the same side of one, and therefore its classification of the survivorship curve as senescent, non-senescent, or negative senescent (sensu Vaupel *et al*. (2004)) does not change. The original discrete-time entropy metric, on the other hand, changes from above one to below one for the type III curve.

Figure 2A and 2C show how the original and the new entropy metrics are correlated for matrix models from COMADRE and COMPADRE, respectively. The sparseness of data in the bottom panels C and D for plants (COMPADRE) are a consequence of the fact that COMPADRE contains largely stage-structured matrices which required a stage-to-age conversion as described in the methods. We excluded models if the demographic quantities such as population growth rate of the converted stage-to-age model differed from the original stage-structured models by more than 5%, which led to many exclusions and therefore a sparser plot for COMPADRE than for COMADRE (bottom two panels versus top two panels in Figure 2). As a consequence of these exclusions, virtually no models from the original datasets analysed in Bernard *et al*. (2020) were left. Therefore we included stage-to-age converted models from COMPADRE outside of the set analysed by Bernard *et al*. (2020), shown in gray in panels C and D.

**Figure 2:**
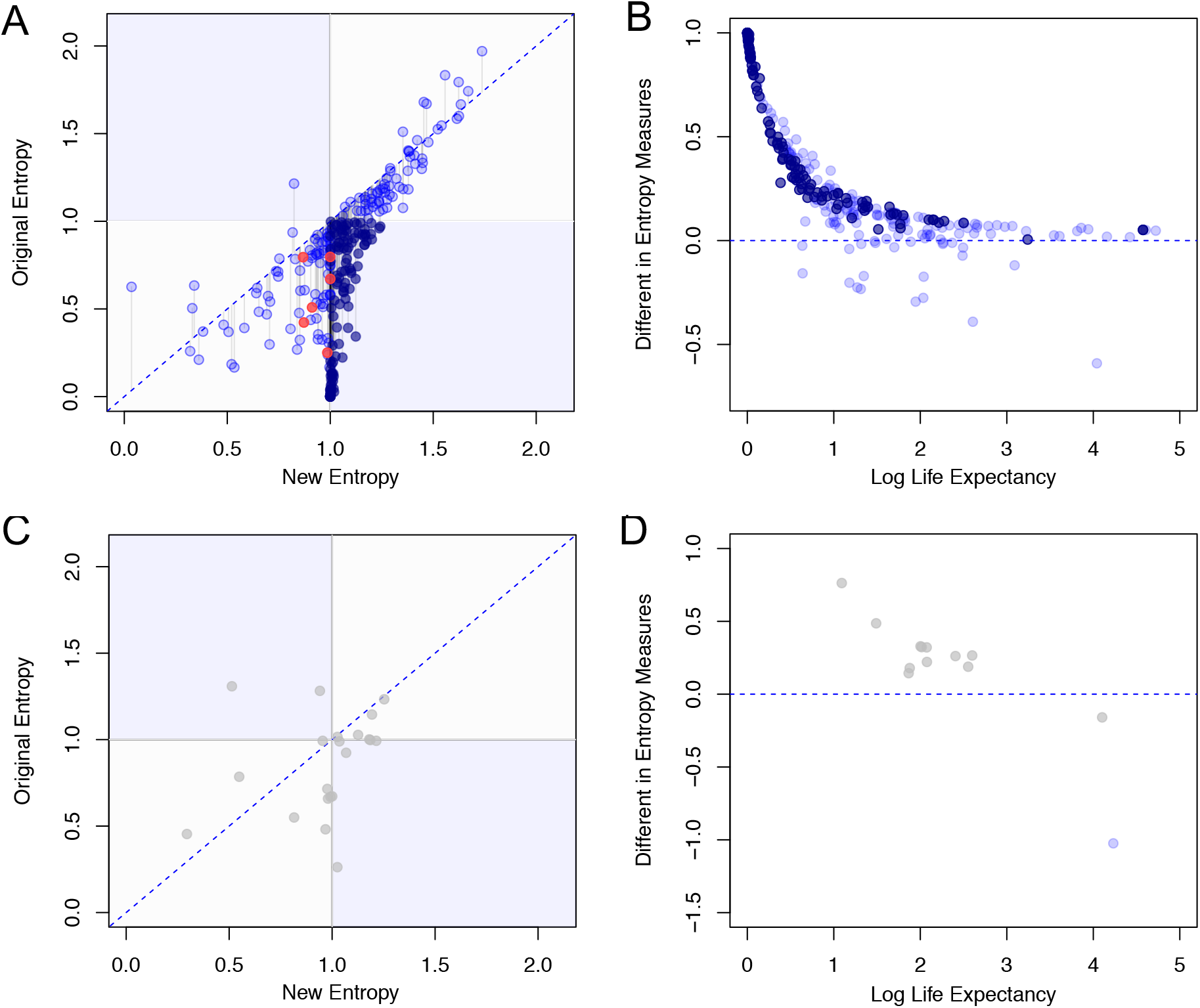
Variation between the original entropy (H_*lx*_) and new entropy (H_*N*_) measures from matrix population models of animals in the COMADRE database (panels A and B), and of plants in the COMPADRE database (panels C and D). A) Comparing Keyfitz’ entropy of animal matrix models from the COMADRE database using the original and new metric. Points in dark blue are matrices where entropy shifted from negative values (positive senescence) to positive values (negative senescence). Matrices converted from stage to age are shown in orange. B) Plot of the difference between the new and the existing metric (H_*N*_ − H_*lx*_) as a function of the life expectancy of animal species from COMADRE. C) Comparing Keyfitz’ entropy of plant matrix models from the COMPADRE database using the original and new metric. Poins in grey are matrix models that were not analysed in Bernard *et al*. (2020). D) Plot of the difference between the new and the existing metric (H_*N*_ − H_*lx*_) as a function of the life expectancy of plant from COMPADRE.

Entropy estimates of animals were correlated between the original discrete-time entropy measure and the new discrete-time entropy formulation, but expressed high variability. The original entropy measure introduced a consistent bias of over-estimating senescence (78.6% of models (264/336) had a greater New Entropy value than Original Entropy value). Bias was weaker at the extremes (colinearly high and low values of entropy) with the strongest overestimation of senescence occurring under the Original Entropy centered at the threshold value where New Entropy = 1 (Figure 2A).

Senescence was strongly underestimated when using the original discrete-time entropy metric (Figure 2A), and also changed sign in a substantial number of cases (two blue surfaces in Figure 2A and C). Sign changed between the two shape measures almost exclusively in the direction of weak negative senescence (small survivorship increase with age) interpreted as positive senescence by the original entropy metric (decreasing survivorship with age; bottom right quadrant of Figure 2A)). Nearly 40% of models (126 of 336) inverted sign between the two entropy measures. Around 34% of the models were more than 0.25 units entropy in error, and 18% of models were more than 0.50 units entropy in error.

The distance between the two entropy metrics was correlated with life expectancy (Figure 2B and D). For data from COMADRE (panel B), the mean error for models with life expectancy < 2 was −0.63, the mean error with life expectancy between 3-5 years was −0.15, and for 5-10 years it was −0.06.

These findings have important implications for the recent and future waves of comparative demographic research evaluating the number of species escaping or undergoing actuarial senescence (e.g., Salguero-Gómez (2017); Beckman *et al*. (2018); Capdevila *et al*. (2020); Bernard *et al*. (2020)) because Figure 2A implies that these studies have likely underestimated the number of species with negligible actuarial senescence using this Keyfitz metric, and Figure 2B implies that this underestimation was especially strong for short-lived species, therefore introducing a spurious correlation between shape and pace.

In Supplementary Materials 1 we show why the original discretisation, *H_lx_*, classifies constant mortality curves as negatively senescent curves, and why it does so more strongly for species with shorter lifespans. We find that for constant mortality *μ*, 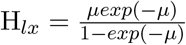. This function is smaller than 1 whenever the mortality rate is nonzero, i.e. when *μ* is larger than 0. Furthermore, 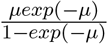 is a decreasing function of *μ* for nonnegative values of *μ* so that the original Keyfitz metric gets closer to zero as the constant mortality gets larger and life expectancy gets shorter.

## 4 Discussion and Conclusion

We have shown that the commonly used time-discretized formula for Keyfitz’ entropy (referred to here as the original entropy measure) does not preserve the relationship between Keyfitz’ entropy and the shape of the survivorship curve that exists for a continuous-time definition of survivorship entropy. That is, constant mortality curves do not yield a Keyfitz’ entropy of one, and life histories with decreasing mortality will not always yield values above one (e.g., see Table 1). Furthermore, the distance between the original and the new Keyfitz’ entropy metric correlates with life expectancy (figure 2B). As a consequence, any correlations obtained between pace and shape of life in previous publications using the existing Keyfitz metric may need to be reevaluated.

We propose a different formula for the discretisation of Keyfitz’ entropy (referred to here as the new entropy measure), based on life disparity and life expectancy in Equation 10. We show in Supplementary Materials 2 that this new formula does preserve the relationship between the shape of the survivorship curve and Keyfitz entropy (that is, *H_N_* > 1 when mortality is a decreasing function of age, *H_N_* < 1 when mortality is an increasing function of age, and *H_N_* = 1 when mortality is constant). However, a major downside of the formula we have proposed is that it is only a measure of the shape of ageing for age-structured survival matrices (Leslie matrices). If the survival and population matrix are stage-structured, then the new entropy measure quantifies whether mortality rate increases or decreases with stage. Stage-to-age conversion methods can offer one way around this limitation to the method.

Due to the constraint of our proposed metric to age-structured matrices, it might in general be better to use one of the many other shape metrics that have been proposed. For example, other life table statistics that have been used to quantify the age-specific decline in survival include Hayley’s median (Hailey, 1874); the age-dependent mortality parameter of mortality distributions (e.g., Gompertz, Weibull, Siler, Logistic, etc.; Ricklefs & Scheuerlein (2002)); the age at the onset of senescence (Jones *et al*., 2008) and the integration of the remaining lifespan and survival function. Wrycza *et al*. (2015) highlight a number of other potential candidates, such as a modified Gini coefficient (reviewing 7 possible metrics), and highlight the value of the entropy as a measure of a statistic of the shape of life (see also Aburto *et al*. (2021) for a recent discussion of measures of lifespan inequality).

Keyfitz derived his continuous-time measure by considering a proportional change in mortality at all ages (section 4.3 in Keyfitz & Caswell (2005)). We did not deduce our formula from this starting point, and instead used existing discrete-time expressions for the numerator and denominator in the continous-time expression derived by Keyfitz. Therefore it remains to be shown whether our new expression for *H*, *H_N_* in equation (10), can also be derived by following Keyfitz’ proof and considering a proportional change in mortality at all ages in a discrete-time model.

## Data accessibility statement

Code used in this paper can be found at https://github.com/Lotte-biology/Keyfitz.

## 1 Supplementary Materials 1: Properties of existing Keyfitz entropy discretisation

### Constant mortality

We start from the definition of Keyfitz entropy given in equation (5) in the main text,

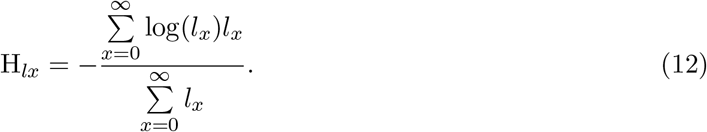

Next we assume mortality is constant, *μ*, such that survival up to age *a* is *l_x_* = exp(−*μx*), and

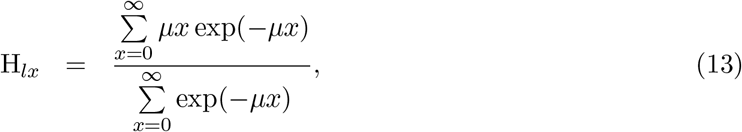

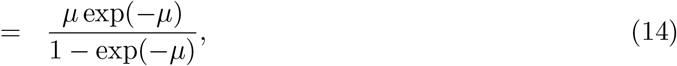

where we used the following two equations for power sums

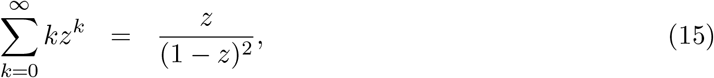

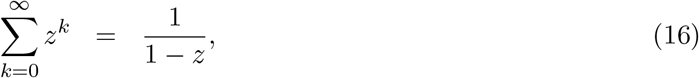

for the numerator and denominator of equation 13, respectively.

In general, H_*lx*_ is therefore not equal to one for constant mortality. Note however, that as we let *μ* get very small, a Taylor expansion of equation 14 shows that it becomes approximately one in the limit of small *μ*. Integrals can be approximated by a series of sums with infinitessimally small step sizes. In the limit of infinitessimally small time steps, survival becomes close to one (or *μ* close to zero), and therefore the sum approximates the continuous time formula well in this limit.

## 2 Supplementary Materials 2: Properties of new Keyfitz entropy formula

### Constant mortality case

We start from the definition of Keyfitz entropy given in equation (11) in the main text,

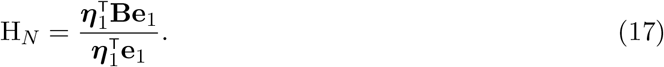

When mortality is constant throughout life, then life expectancy is the same for each age. That is, 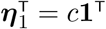 for some constant *c*. Equation (17) then becomes

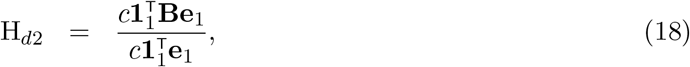

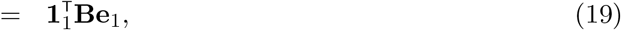

since 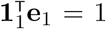. The matrix **B** contains the distribution of age at death, or put differently, each entry contains the probability of dying at that age. Since death is inevitable, the columns of **B** sum to one. That is, for a constant mortality rate

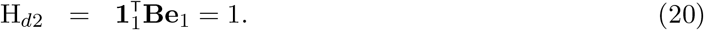

### Increasing mortality rate with age

If mortality rate strictly increases with age, then life expectancy decreases with age, so the entries of 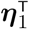 get smaller as you go to higher ages. That is, the first entry of 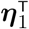, which is the denominator of H_*d*2_, is larger than all other entries of 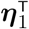. The numerator of H_*d*2_ is a weighted sum of the entries of 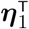, where the weights add to one. Given that the first entry of 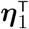 is larger than all other entries, this implies that

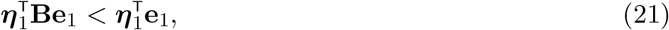

which in turn implies that *H_N_* < 1.

Now we will briefly elaborate why the inequality above is true if the entries of 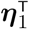 get smaller as you go to higher ages. Lets call the vector with the distribution of each at death for a newborn individual, **Be**_1_ = **p**. We can then write equation (17) as

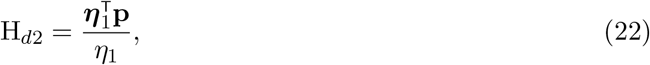

where *η*_1_ is the first entry of the vector 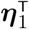. Written out in terms of its components this is equal to

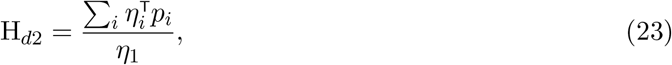

For the case of increasing mortality rates, we know that *η*_1_ > *η_i_* for all *i* > 1. Therefore,

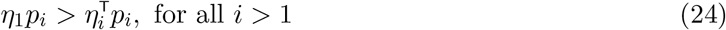

which implies that

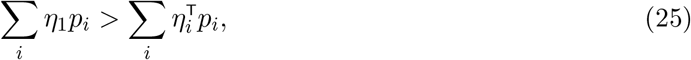

and therefore that

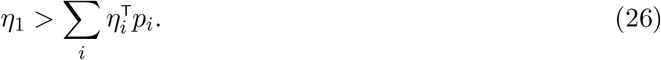

### Decreasing mortality rate with age

If mortality rate strictly decreases with age, then life expectancy increases with age, so the entries of 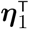 get bigger as you go to higher ages. That is, the first entry of 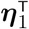, which is the denominator of H_*d*2_, is smallest of the entries of 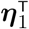. The numerator of H_*d*2_ is a weighted sum of the entries of 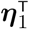, where the weights add to one. Given that the first entry of 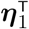 is smaller than all other entries, this implies that

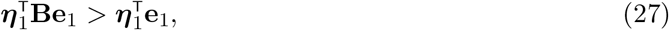

which in turn implies that *H_N_* > 1.

